# Coupling mathematical modeling with a novel human intestinal stem cell system to understand feedback regulation during planar cell polarity

**DOI:** 10.1101/2025.11.05.686808

**Authors:** Keith A. Breau, Emma G. Dunahey, Henry V. Fosnocht, Scott T. Magness, Timothy C Elston

**Author notes:** These authors contributed equally to this work (Scott T. Magness, Timothy C Elston).

## Abstract

The Planar Cell Polarity (PCP) complex regulates many diverse phenotypes. While recent literature has elucidated key mechanisms underlying PCP, a mechanistic understanding of how these components function as a system to drive polarity is lacking. Here, we develop a comprehensive multicellular mathematical model centered around key PCP phosphorylation events, directly simulating the protein interactions that drive PCP. Our model confirms key PCP phenotypes, including robust single-junction asymmetry and multicellular polarity alignment in the absence of extrinsic signals. It predicts unique roles for the two known positive feedback mechanisms and predicts that VANGL-mediated DVL phosphorylation may be an underappreciated negative feedback mechanism. To test model predictions, we employ transgenic primary human intestinal epithelium cultured on biomimetic planar-crypt microarrays (PCMs) as a new platform for studying PCP. Together, our model provides novel insights into the mechanisms that regulate PCP, while our experimental results highlight an unappreciated role for PCP in intestinal biology.

## Introduction

Cellular polarity, the asymmetric organization of cellular proteins and structures, is a fundamental cell process necessary for complex tissue architecture and function. In contrast to apical-basal polarity, which describes sub-cellular asymmetry along the axis perpendicular to the tissue surface (i.e. “top” vs “bottom”), planar cell polarity (PCP) describes the sub-cellular asymmetry of proteins within the plane of a tissue or migrating cell collective (i.e. “front” vs “back”)^1,2^. This coordinated alignment, often mediated by the so-called “core PCP complex”, is crucial across diverse biological contexts, including intestinal patterning^3^, kidney and liver development^4,5^, convergent extension and neural tube closure during gastrulation^6^, alignment of stereocilia bundles required for hearing ^7,8^, the directional beating of respiratory cilia^9^, neural synapse formation and maintenance^10^, hair follicle patterning^11,12^, and angiogenesis^13^. Consequently, disruptions in PCP signaling are implicated in regulating a range of human pathologies, such as neural tube defects^14^, irritable bowel disease (IBD)^15^, Hirschsprung disease^16^, and the progression and metastasis of cancer^17,18^. Thus, understanding the molecular mechanisms governing PCP is key to both deciphering fundamental principles of tissue organization and understanding diverse disease states.

Collective cell migration, or the coordinated directional movement of cell collectives, is a process critical for development, tissue maintenance, and repair^19^ and is often regulated by PCP mechanisms^20,21^. The intestinal epithelium provides two striking examples of such collective behaviors. First, during normal homeostasis, new epithelial cells are constantly born in the stem cell rich crypts, undergo a week-long collective migration to the villus tips, then undergo apoptosis, a life cycle that drives constant tissue renewal^22,23^. Second, following injury, the epithelium undergoes a dramatic coordinated migration process known colloquially as restitution, where cells adjacent to the wound flatten and rapidly migrate *en masse* over the wound bed to reseal the barrier^24,25^. Despite these dramatic phenotypes, how cell polarity is coordinated to orchestrate these events is largely unknown^23^, highlighting this as an important area for further study.

Over the past two decades, considerable progress has been made in elucidating the feedback mechanisms that govern the core PCP complex. Central to the complex is a CELSR-family protein (known as Flamingo or Stan in other model systems), an atypical adhesion-type GPCR that homodimerizes with a CELSR protein in adjacent cells, acting as an essential scaffold for the PCP complex^26–28^. A frizzled-family protein (FZD) then binds to one side of this homodimer to form an initial asymmetry, a key initiating step that stabilizes FZD membrane localization^29,30^. For clarity, we refer to the side of the membrane junction containing the FZD:CELSR complex as the *cis* membrane. In the opposing cell (i.e., the *trans* membrane), this allows for the binding of a VanGogh-family protein (VANGL) to the complex^30^ which in combination with a Prickle-family protein (PRICKLE) drives the clustering of additional VANGL on the opposite membrane^31,32^. Concurrently, the CELSR:FZD complex recruits CK1-family kinases to the *cis* membrane to phosphorylate both VANGL and DVL^33,34^. DVL phosphorylation stabilizes its membrane localization, facilitates multimerization, and increases its binding affinity for FZD^35,36^. By contrast, VANGL phosphorylation on the *cis* membrane reduces its ability to bind or cluster around for oppositely oriented complexes (ie. CELSR-FZD complexes accumulating on the *trans* membrane)^37^. Interestingly, recent evidence suggests that VANGL may also drive the phosphorylation of DVL in the absence of FZD^18,34,38^, a negative feedback mechanism whose role in PCP biology is largely unexplored.

Beyond these core clustering and phosphorylation mechanisms, several other secondary regulatory mechanisms also contribute to the PCP reaction network. PRICKLE for example, promotes the endosomal recycling of most other PCP proteins, including FZD, VANGL, CELSR, and DVL^39^, while VANGL helps recruit both DVL and PRICKLE to the membrane^40,41^. Interestingly, VANGL phosphorylation has a secondary role in this context, both protecting it from ubiquitin-mediated degradation and reducing its affinity for PRICKLE, leading to a general increase in its membrane stability^42,43^. While not an exhaustive list, these mechanisms underscore the complexity of the PCP feedback systems and highlight the need for systems-level approaches to understand how these mechanisms work together to achieve tissue-scale phenotypes.

Mathematical modeling has played a significant role in complementing experimental studies of planar cell polarity by helping connect experimental observations to tissue-scale phenotypes^44^. A key example of this is the Amonlirdviman 2005 model^45^, which was the first to demonstrate that multi-cellular phenotypes can arise from purely local interactions. While previous models have successfully captured certain key PCP phenotypes, their frameworks are generally based on an abstract or simplified view of the core PCP machinery^45–52^. They also often require an assumed extrinsic diffusible gradient as a biasing signal to generate robust polarity, a requirement not currently supported by in vivo experimentation^53,54^. With PCP defects increasingly tied to human disease, a robust and fully mechanistic model of PCP would be a powerful tool for investigating new therapeutic interventions.

Here, we develop a comprehensive mathematical model of the core planar cell polarity complex, focused on the phosphorylation events governing PCP and including all well-supported component interactions of the PCP signaling network. We perform a broad parameter search to find parameters that generate single-junction asymmetry in the absence of biasing signals, exploring how VANGL-mediated negative feedback and biological noise impact model behavior. We then expand the reaction network to a two-dimensional tissue model, exploring the mechanisms that govern multicellular polarity alignment and domineering non-autonomy phenotypes. Finally, we test our model predictions in vivo, using a primary human intestinal VANGL2 overexpression cell line grown on biomimetic planar crypt-microarray scaffolds to demonstrate a role for PCP in the intestinal epithelium and confirm our model’s predictions. Taken together, our results provide important insight into the PCP reaction network, connecting component planar cell polarity interactions to tissue scale phenotypes and providing a novel foundation to understanding the complex role of PCP in human development and disease.

## Results

### Conceptual framework for modeling the PCP reaction network

In the last decade, numerous advances have been made in our understanding of the mechanisms governing Planar Cell Polarity, but no mechanistic model has been produced that incorporates these mechanisms. At the core of our modeling are six proteins, CELSR, FZD, VANGL, DVL, ANKRD6, and PRICKLE in humans (**Fig 1A**), which work together to form the core positive feedback loop that we break down into several key conceptual steps (**Fig 1B**):

**(i)** CELSR family proteins (also known as Stan, StarryNight, or Flamingo in other model systems) are adhesion-mediated G-protein coupled receptors (GPCR) that form homodimers between the membranes of adjacent cells to form a scaffold upon which PCP complexes form.
**(ii)** FZD-family receptors, atypical GPCRs well-known for their central role in WNT signaling, bind to these homodimers to form a stable trimeric complex. Binding of FZD to CELSR disfavors the binding of a second FZD on the opposing membrane. DVL binding to FZD helps stabilize these FZD-CELSR complexes. This asymmetry is a key component of PCP. Here, we use cis to refer to the side of the PCP complex containing the CELSR/FZD heterodimer and trans to refer to the side of the complex in the adjacent cell.
**(iii)** The binding of FZD to CELSR homodimers enables the recruitment of VANGL proteins to the trans membrane. VANGL-family proteins (known as VanGogh or Stbm in other model systems), are four-pass integral membrane proteins that, together with their key binding partner PRICKLE, form the major trans component of PCP complexes. VANGL also promotes recruitment of both DVL and PRICKLE from the cytoplasmic compartment to the membrane.
**(iv)** While the binding of VANGL alone provides partial stability to nascent PCP complexes, the additional binding of PRICKLE to the trans complex facilitates a clustering of VANGL molecules on the trans membrane, further stabilizing nascent PCP complexes. By contrast the binding of PRICKLE to any proteins not in a PCP complex promotes their endocytosis.
**(v)** The binding of FZD to CELSR homodimers enables the recruitment of CK1-family kinases to phosphorylate both DVL and VANGL on the cis membrane. As the regulation of CK1 is not currently believed to be a significant component of the PCP feedback network it was not directly modeled, and its concentration was assumed to be non-limiting. Phosphorylation of DVL-family proteins, also well known as a central player in both canonical and non-canonical WNT signaling, enhances their membrane stability both directly, through increased membrane affinity, and through secondary dimerization. This phosphorylation is modeled as ANKRD6 dependent, as ANKRD6 is a known scaffolding protein of CK1, and is inhibited by the competitive binding of PRICKLE to DVL on the cis membrane. Phospho-DVL in turn diffuses and stabilizes the cis side of nascent PCP complexes, completing the core positive feedback loop. By contrast, phosphorylation of VANGL on the cis membrane inhibits both its ability to bind FZD-CELSR complexes on the trans membrane (opposite orientation complexes) and its ability to be recruited into corresponding opposite orientation VANGL clusters. Thus, VANGL phosphorylation provides the key inhibitory mechanism needed to disrupt the opposite orientation positive feedback loop and achieve an asymmetric distribution of PCP complexes across the junction.

**Figure 1.**
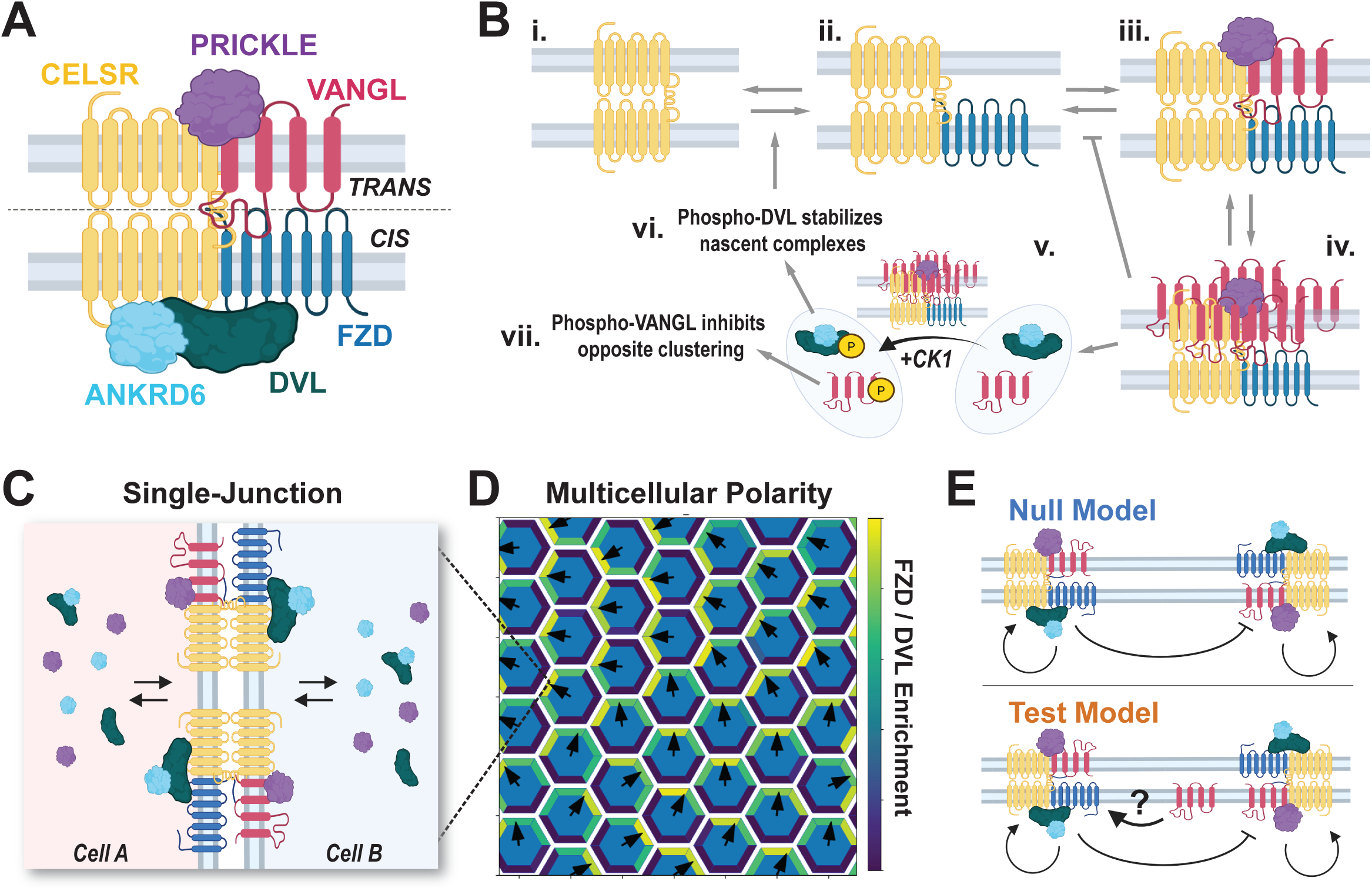
Overview of Modeling Approach **(A)** Proteins included in the model. Cytoplasmic proteins DVL, PRICKLE, and ANKRD6 shuttle between cytoplasmic and membrane-associated states. **(B)** Conceptual framework for the core PCP positive feedback loop, showing: i) intercellular CELSR dimerization, ii) recruitment of on one membrane, iii) recruitment of VANGL and PRICKLE to the opposite membrane, iv) formation of a VANGL cluster, v) Stabilized FZD phosphorylation complex, phosphorylating DVL and VANGL. Phospho-DVL stabilizes CELSR-FZD complexes forming a positive feedback loop and phosphorylated VANGL is inhibited from interacting CELSR, promoting asymmetric complex formation across the junction. **(C)** Graphical representation of the “single-junction” modeling approach, where each cell is represented by a cytoplasmic compartment and a single shared membrane junction. **(D)** Example output from the “multicellular” modeling approach, where each cell is modeled as a cytoplasmic compartment surrounded by six membrane compartments, each part of a membrane junction. **(E)** Graphical representation of the key difference between the two reaction schemes tested. In the “Test” model, the build-up of VANGL caused by VANGL clustering helps stabilize nascent FZD:CELSR complexes independently recruiting CK1 to phosphorylate DVL as a form of negative feedback.

### Mechanistic Modeling Approach

To model this system, we employ a compartmental ODE approach, explicitly modeling the six core PCP proteins’ synthesis, interactions, and degradation using mass-action kinetics. We first perform a broad parameter search using “single-junction” simulations, where each cell is modeled as a cytoplasmic compartment and a single membrane interface (**Fig 1C**), to identify and analyze parameter sets that generate an asymmetric distribution of PCP complexes across the junction. In mathematical terms, this behavior is referred to as bistability. The membrane that becomes the cis membrane is determined randomly by biological noise included in the system. We then use those parameters to analyze behavior in a two-dimensional multi-cellular system, where each cell is represented as a well-mixed cytoplasmic compartment and six membrane junction compartments (**Fig 1D**). Membrane proteins not in PCP complexes diffuse laterally between membrane compartments, and cytoplasmic proteins also come on and off the membrane. The full list of reactions and parameters can be found in **Supplemental Figs 1 and 2**.

**Figure 2.**
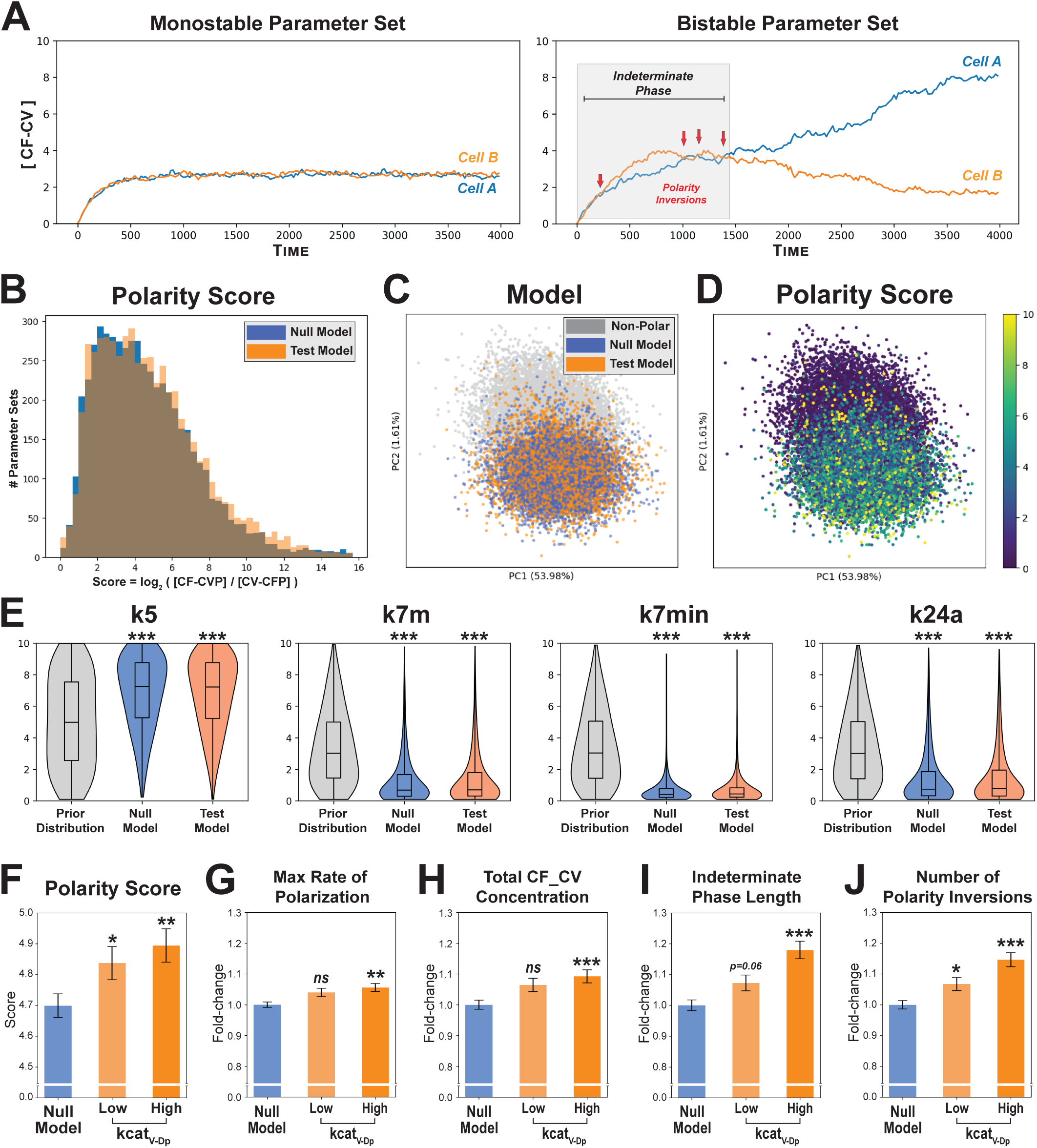
VANGL clustering drives single-junction bistability **(A)** Representative simulations from single-junction modeling, showing the concentration of cis-oriented (CELSR:FZD) complexes in each cell over time. Bistable parameter sets typically display an initial indeterminate phase, where the polarity may invert several times (red arrows) before final polarity is determined. **(B)** Every parameter set demonstrating bistability, scored based on the log fold-change of the ratio of cis-oriented vs trans-oriented complexes. **(C)** PCA analysis of parameter space, including all bistable parameter sets and an equal number of monostable parameter sets. **(D)** PCA representation from (C), overlaid with the computed polarity scores from (B). **(E)** Most significant parameters identified to facilitate bistability. k5 - rate of VANGL binding to the PCP complex; k7m and k7min - effect of VANGL clustering on complex stability; k24a - dissociation rate of non-phosphorylated VANGL from membrane clusters. Box plots show median and interquartile range. **(F-G)** Metrics comparing the effect of no negative feedback (Null Model) versus increasing feedback strength (kcat_VDp_, Low vs. High) across all bistable parameter sets. Data plotted as mean +/- SEM. Wilcoxon rank-sum test. * p < 0.05. ** p < 0.01. *** p < 0.001.

Importantly, it is currently unclear whether any negative feedback loops exist to regulate the system and if so, what effect this feedback would have on complex behavior. One likely candidate feedback loop is the VANGL-dependent phosphorylation of DVL at the membrane, independent of FZD. Such feedback regulation has significant support in the literature, specifically: 1) DVL is phosphorylated even when it can’t bind FZD^55^, 2) VANGL, CK1, DVL form a stable trimeric complex by themselves^34^, and 3) VANGL overexpression or knockdown directly correlated to phospho-DVL levels^38^. We reason that VANGL-dependent phosphorylation of DVL likely forms a negative feedback loop because VANGL clustering on the trans membrane increases the overall level of VANGL (both clustered and free). This free VANGL in turn increases the amount of phosphorylated DVL, promoting the formation of FZD-CELSR heterodimers on the trans membrane and decreasing the asymmetry of the junction. To test how this feedback impacts model behavior, we compare a Null model (no negative feedback) with a Test model in which free VANGL stabilizes FZD-CELSR complexes via VANGL-dependent DVL phosphorylation (**Fig 1E**).

### Both models demonstrate robust bistability in the absence of biasing signals

To determine whether our models were capable of spontaneous polarity establishment, we generated five million random parameter sets for each model. To balance a wide parameter space with computational feasibility, we used a 100-fold range for most parameter values, with smaller ranges for synthesis and degradation rates and a set of biologically derived constraints to help reduce the overall parameter space (**Supp Table 1 and Methods**). Simulations were iterated forward in time using the Euler method until >99% of simulations had reached steady state, then analyzed for bistability (i.e. asymmetric accumulation of PCP complexes). All simulations were conducted with a small amount of Gaussian white noise added to the synthesis rates of all proteins and without any biasing signal (see Methods).

Interestingly, many parameter sets demonstrated bistability (*n*=5411 and *n*=5637 for Null and Test models respectively), often following an initial indeterminate phase where the polarity may invert several times (**Fig 2A**). The scale of polarity varied between parameter sets, with a mean log-2 fold-change in complex enrichment of ∼4.8 (∼28-fold enrichment, **Fig 2B**). To determine whether polarity establishment was following a single shared mechanism, principal component analysis of parameter space was performed, including all polarizing parameter sets from both models along with an equal number of randomly selected non-polarizing controls. Our results demonstrate only a single polarizing basin within parameter space, suggesting that all polarizing parameter sets are achieving bistability using a single broadly shared mechanism (**Fig 2C,D**).

### VANGL-mediated positive feedback drives single-membrane polarity

To determine which parameters were most critical to polarity establishment, we next performed rank sum statistical testing comparing the initial distribution of each parameter to its distribution within the polarizing parameter sets. Interestingly, the top 11 most significant parameter values were all related to VANGL-mediated complex stabilization, with the most significant being a high rate of VANGL binding to the PCP complex (<u>k5</u>), a strong effect of VANGL clustering on complex stability (<u>k7m</u> and <u>k7min</u>), and a low rate of non-phosphorylated VANGL dissociation from membrane clusters (<u>k24a</u>, **Fig 2E**). Together, these data support a strong central role for VANGL in driving membrane bistability at the single-junction level.

### Negative feedback delays polarity establishment, but slightly enhances final polarity strength

To better understand how negative feedback via VANGL-mediated DVL phosphorylation impacts polarity establishment, we split the pool of polarized Test model parameter sets in half based on the phosphorylation rate (kcat_V-<u>Dp</u>) and tested for significant differences in key polarity metrics versus the Null model. While a slight increase was observed for the Test model in the mean polarity score, max polarization rate, and final PCP complex concentration (**Fig 2F-H**), the most significant differences were an increased average length of the indeterminate phase (∼18% increase, **Fig 2I**), coupled with a larger number of polarity inversions during this phase (∼15% increase, **Fig 2J**).

### Multicell modeling demonstrates robust polarity alignment in the absence of gradients

The question of whether the PCP reaction network can generate polarity alignment over a population of cells in the absence of external signals such as WNT ligand gradients is an open debate. To investigate this question, we extended our two-membrane model to multicell models. In these models, cells are treated as hexagons with each edge of the cell representing a membrane compartment surrounding a shared cytoplasm (**Fig 1D**). Using bistable parameter sets found in the two-cell model, we ran multicell simulations for both the Null and Test models. Strikingly, many parameter sets in both models generated robust multicell alignment in the complete absence of exogenous biasing signals (**Fig 3A**).

**Figure 3.**
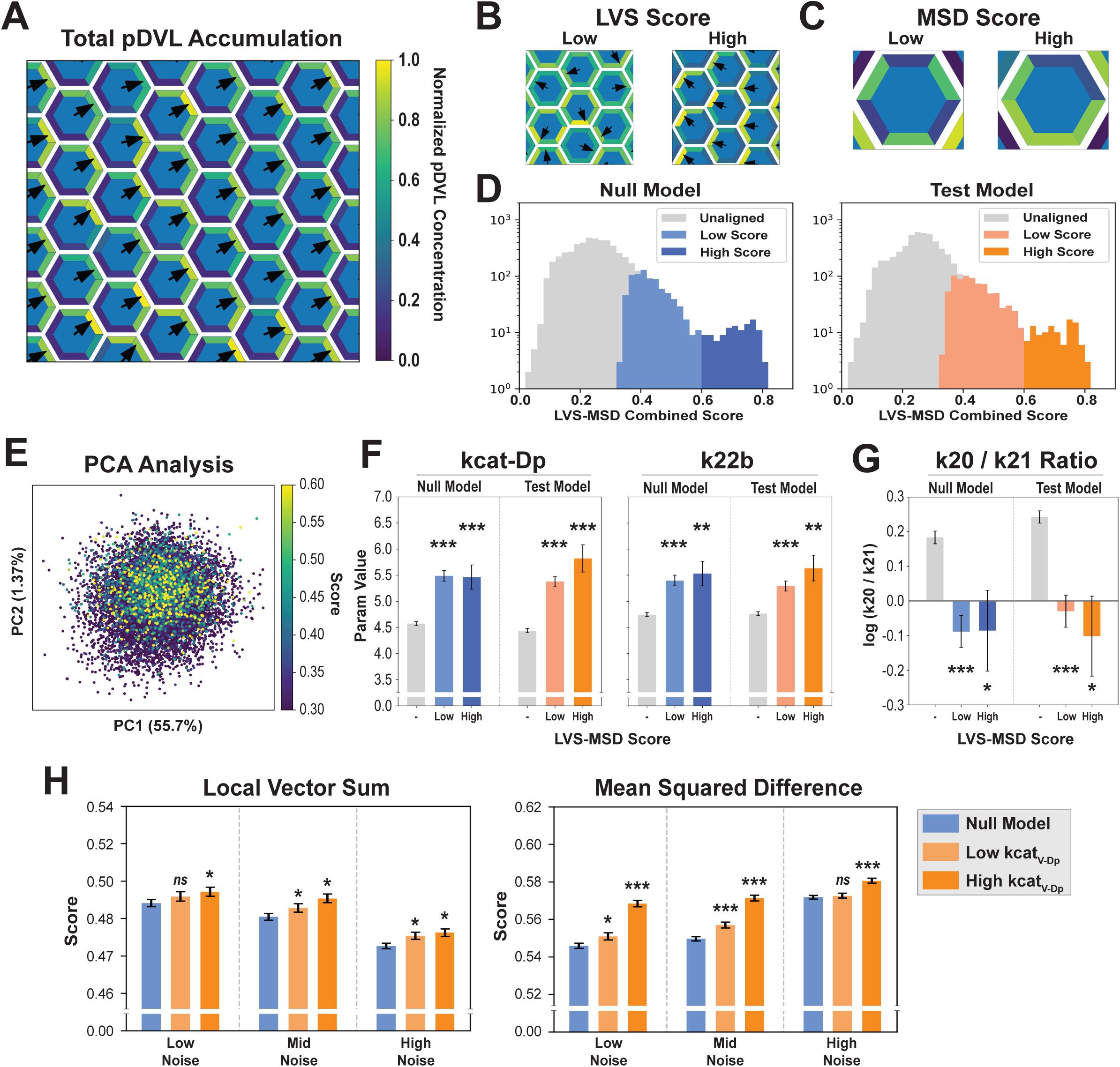
Regulated DVL phosphorylation mediates polarity alignment **(A)** Representative multicellular simulation showing robust polarity alignment in the absence of any biasing signal. **(B,C)** Example images demonstrating the two multicellular scoring metrics employed. LVS measures inter-cellular alignment while MSD measures intra-cellular alignment. **(D)** Stacked histograms of the combined LVS*MSD score for every parameter set tested, comparing the score distributions of parameter sets designated “Unaligned”, “Low Score” or “High Score” for each model. **(E)** PCA analysis of all parameter sets used in multicellular simulations, overlaid with the LVS*MSD score for each. **(F)** Average parameter value in Unaligned, Low Score, and High Score parameter sets from (D). kcat-Dp – rate of DVL phosphorylation by FZD:CELSR complexes. k22b – cytoplasmic DVL dephosphorylation rate. Mean +/- SEM. **(G)** Ratio of phospho-DVL dimerization (k20) and de-dimerization (k21) rates. **(H)** Alignment metrics comparing the effect of no negative feedback (Null Model) versus increasing feedback strength (kcat_VDp_, Low vs. High) across all simulated parameter sets. Data plotted as mean +/- SEM. Wilcoxon rank-sum test. * p < 0.05. ** p < 0.01. *** p < 0.001.

### Metrics to quantify polarity alignment

To quantitatively compare the alignment of different parameter sets, we developed two metrics. The first metric, called Local Vector Sum (LVS), measures how well the total polarity vector (the vector sum of cis-oriented complexes in a cell) of a given cell is aligned to the total polarity vectors of its six local neighbors (i.e <u>inter</u>-cellular alignment, **Fig 3B**). The LVS is computed as the ratio of the actual vector sum magnitude of the seven-cell neighborhood over the theoretical maximum vector sum magnitude if all seven neighbors were perfectly aligned. This is computed for every cell in the simulation, and the mean value is the LVS score for that parameter set.

The second metric, called Mean Squared Difference, measures how smoothly distributed PCP complexes are within each individual cell (i.e. <u>intra</u>-cellular alignment, **Fig 3C**). First the concentrations of cis-oriented complexes in each cell are normalized, such that the highest-concentration membrane in that cell is 1 and the lowest is 0. MSD is then computed for each membrane compartment 𝑋_𝑖_ as: 1 − (𝑋_𝑖_ − (𝑋_𝑖−1_ + 𝑋_𝑖+1_⁄2))^2^, and averaged over the field of cells.

To be a good candidate, a parameter set should produce high values of both measures. Gaussian mixed modeling with two components was performed to identify a minimum threshold for each metric, and parameter sets below this threshold for either metric were discarded. Interestingly, combining both scores into a single metric by forming their product (LVS*MSD) revealed clear populations of well-aligned simulations with a combined LVS*MSD score of >0.6 in both the Null and Test models (**Fig 3D**). To better understand what parameters drive this level of alignment, parameter sets above this threshold were designated as High Score and sets below this threshold, but which passed the filtering for each individual metric, were designated as Low Score for further analysis.

### The DVL-mediated positive feedback loop facilitates cellular alignment

To determine which parameters had a significant role in facilitating polarity alignment, we first performed PCA analysis of parameter space, again finding that well-aligned models shared a single large basin within parameter space and supporting a common alignment mechanism (**Fig 3E**). We then performed rank sum tests comparing the initial parameter distribution to the Low and High scoring parameter sets. In contrast to the role of VANGL in driving initial membrane polarity (**Fig 2E**), this analysis revealed a significant role for DVL phosphorylation in mediating polarity alignment. High scoring parameter sets demonstrated both significantly increased mean rate of DVL phosphorylation at the membrane (<u>kcat-Dp</u>) and increased cytoplasmic de-phosphorylation (<u>k22b</u>), showing that turnover of phospho-DVL is an important component of PCP in mediating local polarity alignment (**Fig 3F**). Interestingly, the ratio of the pDVL dimerization and de-dimerization rates (k20 / k21), was also significantly reduced in high scoring models (**Fig 3G**), showing that excessive trapping of pDVL at the membrane may be deleterious to robust alignment.

### Negative feedback enhances intra-cellular polarity alignment

Our initial investigations in the single-junction system showed that the Null model demonstrated a longer indeterminate phase length, enabling more polarity inversions before final polarity determination (**Fig 2I,J**). Therefore, we wondered if negative feedback might allow the system more time to integrate signals from neighboring cells, improving overall inter- or intra-cellular alignment. Alternatively, we questioned whether negative feedback might act as a noise filter, helping facilitate robust alignment in more noisy contexts. To test these possibilities, we re-ran our previous multicell simulations for both models using 10-fold or 100-fold increased noise. To help visualize the impact of VANGL-mediated DVL phosphorylation, parameter sets in the Test model were separated into upper and lower halves based on their value for the catalytic rate kcat_V-<u>Dp</u>.

Analysis of the LVS scores found only a minimal increase in polarity alignment regardless of noise level, indicating that negative feedback does not have a strong role in promoting inter-cellular polarity alignment or in noise filtering (**Fig 3H**). By contrast, the Test model demonstrated significantly improved intra-cellular alignment (MSD) across all noise levels, indicating that negative feedback could play an important role in preventing excessive complex accumulation at any given membrane. Interestingly, MSD score also increased with noise, likely due to a similar effect of reducing excessive complex accumulation. These results further support the notion that VANGL-mediated DVL phosphorylation has beneficial impacts to PCP network alignment.

### The role of WNT gradients in directing PCP

While the necessity for WNT gradients to drive PCP in certain biological contexts is still unclear^53,54^, their impact in other contexts has been well established^18,38,56^. To assess how WNT ligands interact with the PCP complex to drive multicellular alignment, we simulated a WNT gradient over a field of cells with periodic boundary conditions. To simulate a WNT gradient we used a WNT concentration profile that was maximum at the center of the domain and then exponentially decayed in all directions (**Fig 4A**). At each cell membrane, the WNT signal was assumed to increase the phosphorylation rate for DVL and VANGL (kcat-Dp and kcat-Vp, respectively), replicating the primary effect of WNT ligands in vivo^38,57^ (**Fig 4B**). All parameter sets demonstrating modest alignment were simulated (**Fig 3D**), then parameter sets demonstrating consistent alignment over the length of the gradient were analyzed further.

**Figure 4.**
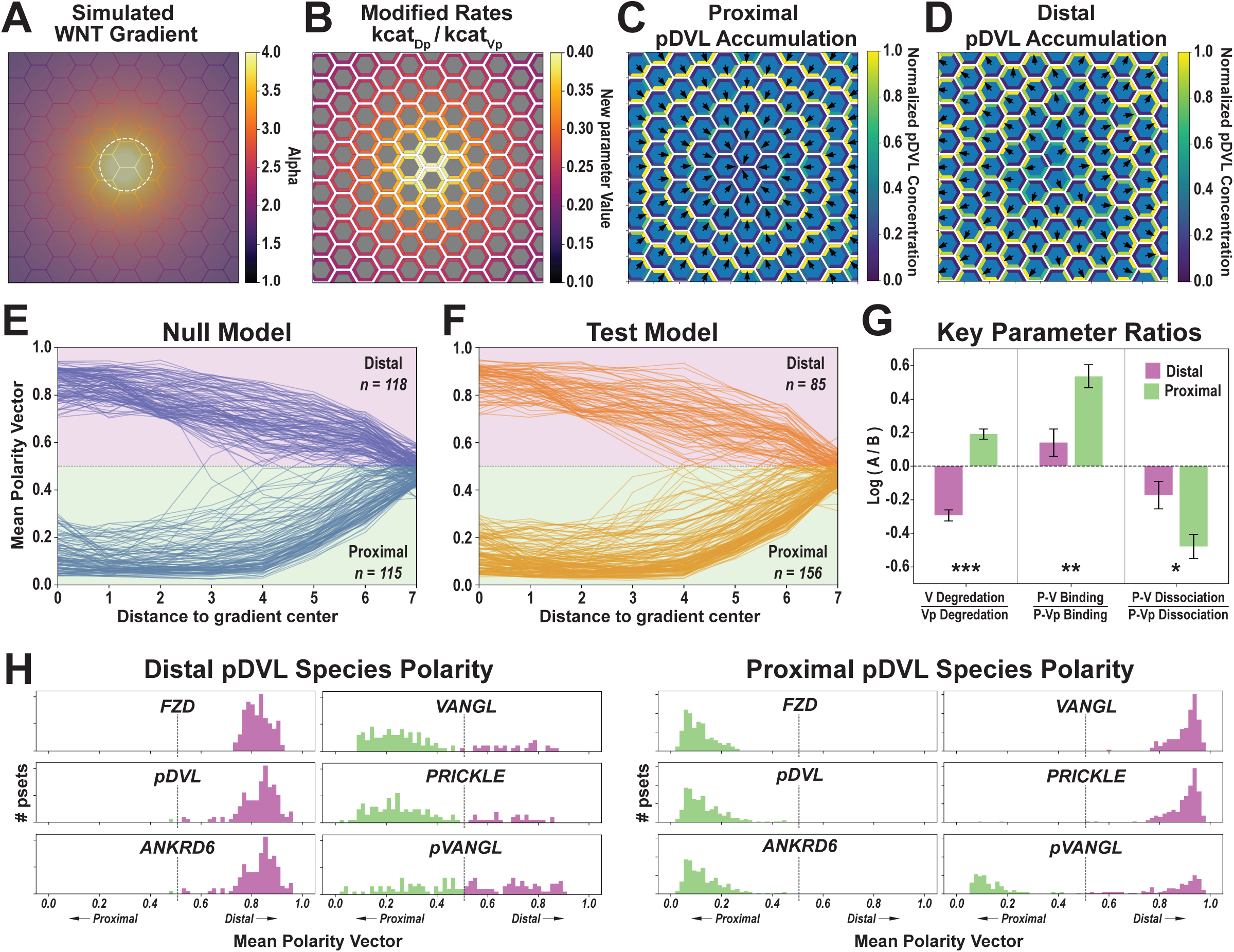
Modeling demonstrates multiple potential responses to a WNT gradient **(A)** Visual representation of the applied WNT gradient, including a central high-WNT region (dashed white circle) outside of which the signal decays exponentially. **(B)** Example simulation, showing how the continuous gradient from (A) is converted into discrete parameter value increases at each membrane compartment. **(C,D)** Representative results from WNT gradient simulations of different parameter sets showing how phospho-DVL can alternatively accumulate either proximally (C) or distally (D) from the WNT source. **(E,F)** Polarity profiles of different parameter sets under a simulated WNT gradient. Each line represents a unique parameter set and indicates whether phospho-DVL is accumulating on the side of the cell proximal or distal to the gradient center. **(G)** Comparison of parameter ratios that are significantly different between proximal-accumulation and distal- accumulation phenotypes. Data plotted as mean +/- SEM. **(H)** Histograms comparing the accumulations of key PCP proteins between proximal-accumulation and distal- accumulation phenotypes. Distal-accumulation simulations show distal accumulation of FZD, phospho-DVL, and ANKRD6 with poorly polarized accumulation of VANGL, PRICKLE, and phospho-VANGL. Proximal-accumulation simulations show overall more robust polarity, with FZD, phospho-DVL, and ANKRD6 accumulating on the proximal cell edge and VANGL and PRICKLE robustly accumulating on the distal edge. Wilcoxon rank-sum test. * p < 0.05. ** p < 0.01. *** p < 0.00

Surprisingly, two opposing phenotypes were observed in response to a WNT gradient. For some parameter sets, pDVL accumulated on the proximal side of the cell relative to the gradient center (**Fig 4C),** while in others it accumulated on the distal side (**Fig 4D**). While at least one instance of proximal accumulation of pDVL has been experimentally observed^58^, WNT ligands are known to be alternatively either attractive^59^ or repulsive^60^ to migratory cells, and our results support that both responses are possible under different biological contexts (see Discussion). Interestingly, the Null model demonstrated roughly equal proportion of these responses (**Fig 4E**), while the Test model is weighted towards proximal pDVL accumulation (**Fig 4F**), suggesting that negative feedback helps promote the proximal pDVL accumulation phenotype.

### The relative recycling rates of VANGL and pVANGL regulate the gradient response phenotype

Given these unexpected findings, we next questioned what parameters influenced the choice between the two opposing phenotypes. It has been previously observed that VANGL phosphorylation has a dual role in regulating its recycling, protecting it from ubiquitin-mediated degradation^43^ while simultaneously reducing its PRICKLE binding affinity^42^ and endosomal internalization^43^. Interestingly, the most significant parameters identified when comparing opposing gradient responses were the VANGL degradation rate and PRICKLE binding affinity. Parameter sets with proximal pDVL accumulation best represented experimentally observed degradation mechanics, with non-phosphorylated VANGL demonstrating increased degradation, increased PRICKLE binding affinity, and decreased PRICKLE dissociation, relative to pVANGL (**Fig 4G**). Furthermore, the degree to which key PCP proteins polarize within the cell is significantly enhanced in the proximal pDVL alignment phenotype compared to the distal phenotype, which often demonstrates poor polarization of VANGL and PRICKLE (**Fig 4H**). Together, these results suggest that regulating the relative degradation and internalization of VANGL and pVANGL is an important component in the PCP response to WNT ligands and predict that disruption of these processes may alter in vivo WNT response phenotypes.

### The Test model predicts a unique domineering non-autonomy phenotype

Domineering non-autonomy is a widely used phenotype for studying PCP, describing a phenomenon where PCP mutant cells have a dominant effect on the polarity orientation of neighboring wild-type cells. Depending on the mutant, this can drive DVL accumulation either proximal (**Fig 5A**) or distal (**Fig 5B**) to the mutant. Both the Null and Test models demonstrated the expected phenotypes for FZD overexpression (distal DVL accumulation), FZD knockout (proximal DVL accumulation), and VANGL knockout (distal DVL accumulation) (**Figs 5C,D**). Unexpectedly however, the Test model demonstrated a unique phenotype in response to VANGL overexpression, with a subset of parameter sets showing distal pDVL accumulation, as opposed to the purely proximal accumulation observed in the Null model (**Fig 5E**). Several parameters were significantly different between these two response profiles (**Fig 5F**), with distal pDVL accumulation parameter sets demonstrating enhanced PCP complex stability in the presence of dimerized pDVL (k4d-low), an increased basal dissociation rate of pVANGL from the complex (k8min-high), a higher DVL- ANKRD6 binding rate (k10-high), and a higher rate of VANGL-driven DVL phosphorylation (kcat-VDp). While these parameters do not highlight a single key mechanism driving this response, this phenotype provides a testable hypothesis to further support a role for VANGL-mediated DVL phosphorylation in PCP.

**Figure 5.**
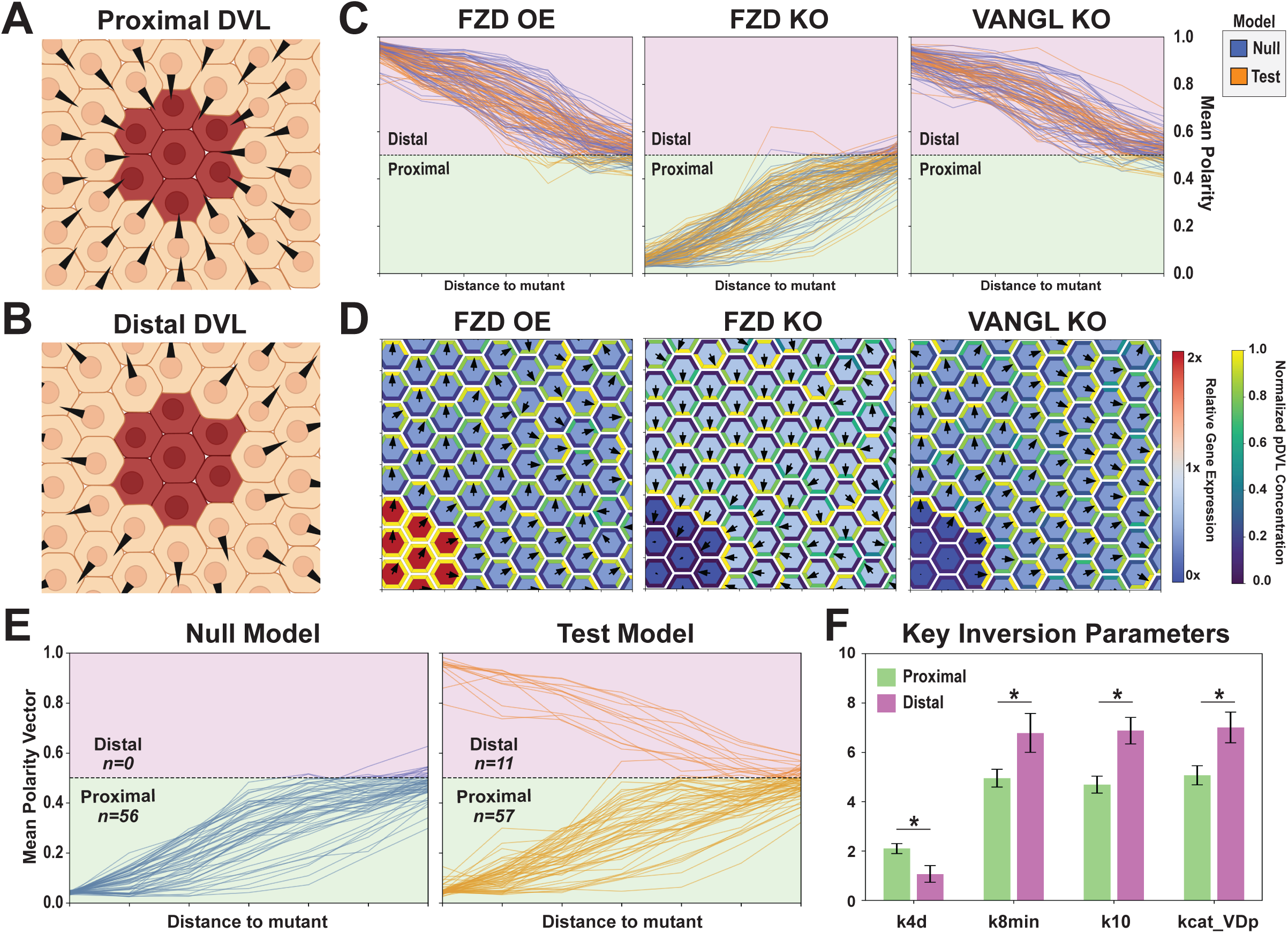
The Test model predicts a unique domineering non-autonomy phenotype **(A,B)** Graphical examples of proximal-DVL (A) and distal-DVL (B) domineering non-autonomy phenotypes. A black triangle in each cell represents the direction of peak DVL membrane activity. **(C)** Polarity profiles of different parameter sets in domineering non-autonomy simulations. Each line represents a unique parameter set and indicates whether phospho-DVL is accumulating on the side of the cell proximal or distal to the gradient center. **(D)** Representative images from each type of domineering non-autonomy simulation in (C). **(E)** Polarity profiles of different parameter sets in VANGL OE simulations. Some parameter sets in the test model demonstrate a unique distal phenotype not observed in the null model. **(F)** Comparison of parameter values between proximal and distal parameter sets from (E). Data plotted as mean +/- SEM. Wilcoxon rank-sum test. * p < 0.05.

### A primary human intestinal epithelial cell platform to test model predictions

To test our predictions in a biological system with strong clinical potential, we next decided to investigate whether the core PCP complex plays a role in the collective migration of primary human intestinal epithelial cells (hIECs). Collective migration is an important component of intestinal epithelial maintenance, observed both during homeostasis^23^ and wound healing^61^, but the mechanisms that coordinate these migratory events are currently poorly understood. To determine whether PCP plays a role in coordinating these events, we developed a hIEC stem cell line to test for domineering non-autonomy phenotypes, using primary human intestinal stem cell culture^62^ and transfection^63^ methods on which we have previously published. Briefly, human intestinal stem cells (hISCs) were isolated from deceased organ donors, expanded in culture, and electroporated using the PiggyBAC™ transposase system to stably integrate one of two DNA constructs (**Fig 6A**)^64^. First, a constitutive H2B-GFP construct, where a nuclear GFP signal enables live tracking of otherwise WT cells (**Fig 6B**). Second, an H2B-mCherry labeled inducible VANGL2 overexpression construct, which selectively overexpresses VANGL2 only in the presence of doxycycline (**Fig 6C,D**).

**Figure 6.**
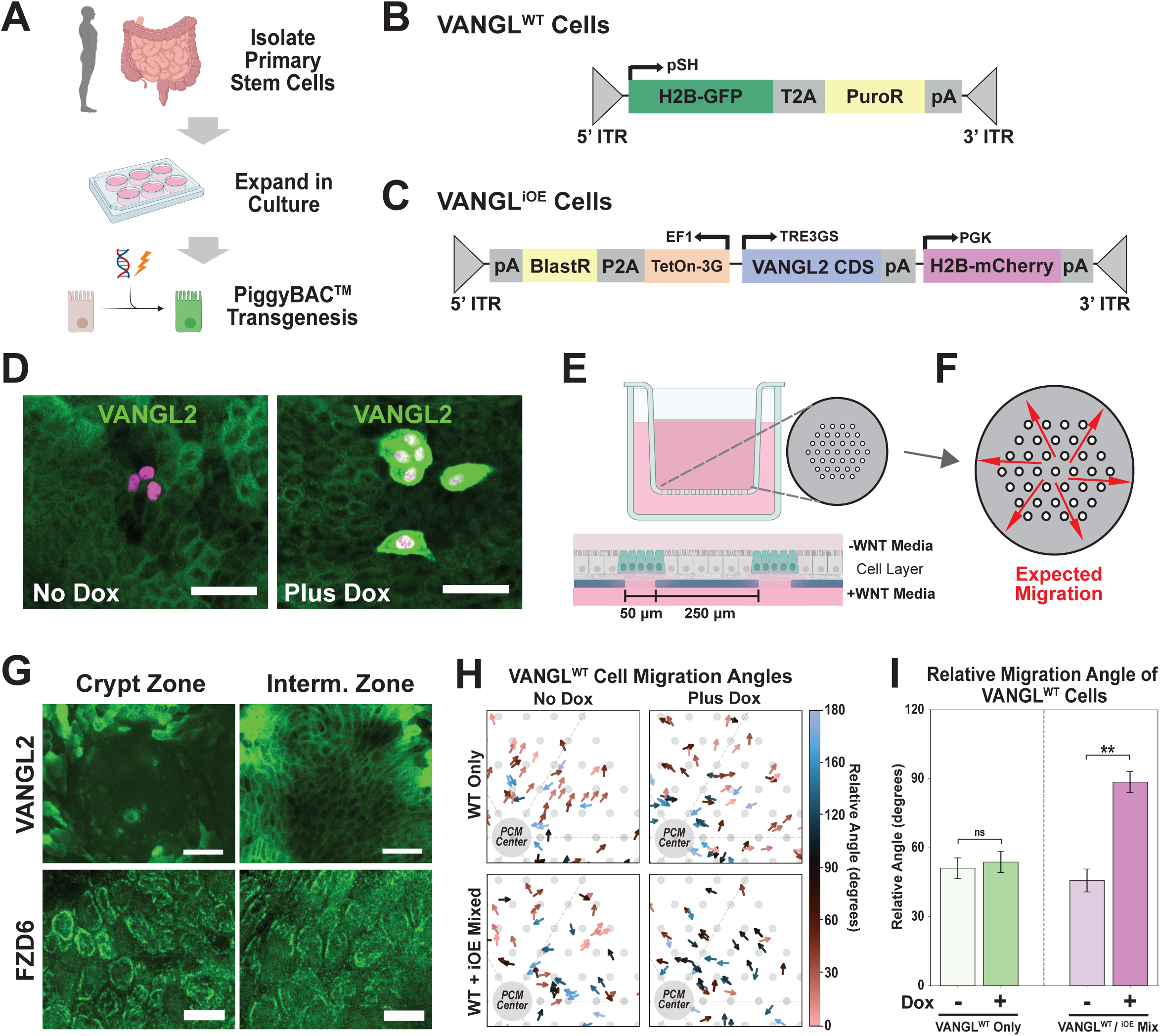
The core PCP complex mediates intestinal epithelial migration **(A)** Graphical overview of workflow for generating transgenic cell lines. **(B)** Schematic of the H2B-GFP constitutive expression plasmid used for tracking cell migration. **(C)** Schematic of the VANGL2 inducible expression plasmid used to test for domineering non-autonomy phenotypes, including a constitutively expressed H2B-mCherry. **(D)** Immunofluorescent VANGL2 staining of VANGL2-iOE cells mixed into WT cultures. In the absence of doxycycline, iOE cells (marked by nuclear mCherry signal) express normal levels of VANGL2 protein. When doxycycline is added to the culture media, iOE cells express high levels of VANGL2. Scale bars 100 um. **(E)** Schematic of planar crypt microarray (PCM) layout. The growth surface of a Transwell is replaced with an impermeable membrane patterned with an array of holes. Culture with canonical WNT ligands in the basal media compartment causes the cell layer to self-pattern in a way that recapitulates in vivo morphology on a 2D surface. **(F)** Overall migratory pattern of differentiated cells after exiting PCM crypt zones. **(G)** Immunofluorescent staining of PCMs. VANGL2 is present at cell junctions in intermediate zones but largely absent from crypt zones. FZD6 is present in both but shows greater cellular asymmetry in intermediate zones. Scale bars – VANGL2, 100 um; FZD6, 25 um. **(H)** Live image cell tracking of VANGL2-WT cells. Each arrow represents the initial and final positions of a tracked cell. Arrows are colored based the angle of migration relative to the PCM center. Grey circles indicate the positions of holes in the PCM membrane. **(I)** Quantification of the migration angles from (H). Addition of VANGL2 OE cells causes a significant shift toward perpendicular migration. Wilcoxon rank-sum test. * p < 0.05. ** p < 0.01. *** p < 0.001

To accommodate live imaging constraints while maximizing biological relevance, these cell lines were combined with a previously published biomimetic platform called planar crypt microarrays (PCMs), which seek to mimic in vivo crypt-villus spatial patterning on a planar surface (**Fig 6E**). In this platform, an impermeable membrane is patterned with an array of holes, adhered to the bottom of a Transwell™ culture insert, and coated with collagen. To mimic the in vivo stem cell niche, stem cell maintenance media rich with canonical WNT ligands is added to the basal media compartment, accessible only to cells within close proximity of a micro-hole. When hISCs are cultured on this platform, stem cell rich crypt regions form over each micro-hole and proliferate continuously^65^. Similar to in vivo biology, new cells born within these crypt regions demonstrate a continuous outward migration into differentiated regions^66^, followed by a general outward migration to the edge of the patterned region (**Fig 6F**).

To assess whether any regions of the PCM platform demonstrated planar polarization of PCP components, we fixed and stained mature PCMs for VANGL2 and FZD6, a Frizzled-family receptor with a well-established role in PCP^67^. Our results show that while VANGL2 is largely absent from the crypt regions, it is present on the membranes of nearly all cells in the intermediate zones (**Fig 6G**). While FZD6 is present in both regions, it shows poor intracellular asymmetry in crypt regions compared to intermediate zones (**Fig 6G**). Together, these observations suggest that PCP mediates cell migration primarily outside of intestinal crypts. Interestingly, previous in vivo observations found that active migration of IECs up the villi only begins following exit from the crypts^23^, further supporting this idea. Thus, further experiments focused only on cells that have already exited the crypt zones.

### Human intestinal epithelium demonstrates domineering non-autonomy migration defects

After exiting crypt zones, hIE cells grown on the PCM platform typically display a net outward displacement toward the edge of the pattern (**Fig 6F**). To determine whether the presence of VANGL2- overexpression mutants has a domineering non-autonomous effect on WT cell migration, VANGL2- WT and VANGL2-iOE cells were combined at an ad hoc 10:1 ratio and grown on PCMs, employing live fluorescent imaging to track cell migration. When WT cells were cultured together with VANGL2- overexpressing neighbors, their migration behavior was significantly altered. Specifically, WT cells displayed a marked shift in the angle of net displacement when co-cultured with VANGL2-OE cells, compared to both WT cultures and uninduced mixed cultures (**Fig 6H-I, Supp Videos 1-4**). Importantly, tracked cells were often many cell diameters away from VANGL2-OE cells, indicating the effects of VANGL overexpression can propagate through intestinal monolayers to disrupt collective migration dynamics. Thus, our results support a general role for PCP in mediating the collective migration of human intestinal epithelial cells.

### A wound-healing assay demonstrates VANGL-OE oriented migration predicted by Test model

While these results support a general role for PCP in hIEC collective migration, the non-uniform nature of migrating cells in this assay made testing our model’s predictions on the directional impact of VANGL2-OE cells inconclusive. We reasoned that changes to migration would be more easily observed in a wound healing assay, as the intestinal epithelium is known for a robust and well-coordinated wound healing response called restitution^25^. As pipette scratches often produce jagged edges on PCMs due to the collagen, we first developed a novel wound assay system. A thin L-shaped strip of PET film was embedded into the collagen off to the side of the micro-hole region, and cells were cultured on top of the collagen layer as previously described. Immediately before live imaging, the protruding tip of the wound strip was grabbed with forceps and pulled up, generating a relatively straight wound in the cell layer (**Fig 7A**). Interestingly, immunofluorescent staining of VANGL2 in WT cells 12 hours post-wounding highlights a significant anterior enrichment in many cells near the wound edge, consistent with a strong degree of PCP complex polarization in these cells (**Fig 7B**).

**Figure 7.**
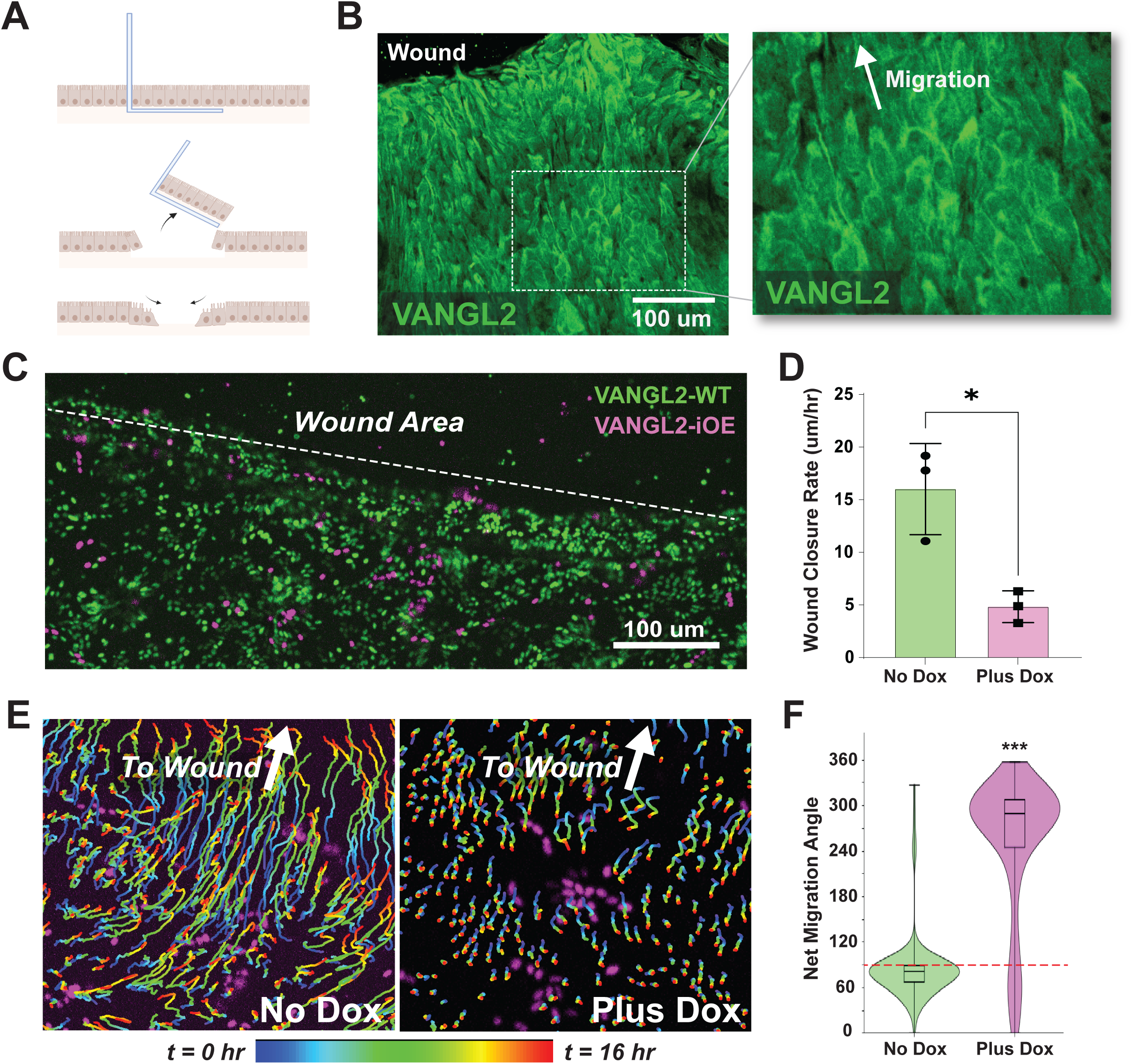
Intestinal epithelial cells migrate towards transgenic VANGL-OE cells **(A)** Graphical overview of wound strip usage. A thin strip of PET film is embedded in the collagen beneath the cell monolayer and torn up to create a wound. **(B)** IF staining of IECs fixed during the wound healing process. Inset shows VANGL2 accumulating in lamellipodia-like structures on the leading edge of migrating cells. **(C)** Representative image of a wound healing assay at t=0, demonstrating a mix of WT (green nuclei) and VANGL2-iOE (magenta nuclei) cells adjacent to a wound. **(D)** Overall wound closure rate averaged along the entire migration front from n=3 experimental replicates. Bars represent mean +/- stdev. **(E)** Example migration tracks of WT cells near patches of VANGL2-OE cells following dox induction demonstrates migration oriented toward OE cells. **(F)** Migration angles of WT cells located between VANGL2-iOE cells and the wound edge. Red dotted line at 90 degrees indicates the direction of the wound. At least four OE patches and 200 WT cells were analyzed for each condition. Wilcoxon rank-sum test. * p < 0.05. ** p < 0.01. *** p < 0.001

To test for domineering non-autonomy defects in wound closure, VANGL2-WT and VANGL2- iOE cells were once again mixed at a 10:1 ratio and cultured on PCMs with embedded wound strips. Test cultures were induced with doxycycline, the wound strip was removed, and live fluorescent imaging employed (**Fig 7C**) to track wound closure. Strikingly, while robust and uniform migration into the wound region was observed in control cultures, a >65% impairment in wound closure rate was observed when VANGL2 overexpressing cells were included in the cell layer (**Fig 7D, Supplemental Video 5-6**). This assay finally allowed us to assess the directional impact of VANGL2 overexpression on nearby regions of the cell layer. The spot tracking feature in the Imaris software suite was used to track WT (GFP+) nuclei in high throughput, turning our live imaging data into cell migration tracks. We frequently observed migration oriented towards VANGL2-OE cells (**Fig 7E**), with WT cells located between a VANGL2-OE patch and the wound edge demonstrating drastic shifts away from wound- oriented migration (**Fig 7F**). As we and others have shown, FZD and pDVL typically accumulate in the rear of collectively migrating cells where they activate RhoA^68^, while VANGL accumulates at the front (**Fig 7B**). Thus, VANGL-OE oriented migration is consistent with the distal pDVL accumulation phenotype only observed in our Test model. Together these results support both a significant role for PCP in intestinal wound healing, and a role for VANGL-mediated negative feedback in PCP regulation.

## Discussion

By combining computational modeling with a new experimental platform for studying planar cell polarity, we discovered unique roles for known feedback mechanisms in the PCP pathway. Our modeling results demonstrate that multicellular polarity alignment does not require extrinsic signals and suggests various ways that negative feedback improves polarity alignment. Our experimental system revealed an important role for PCP in mediating intestinal epithelial migration and confirmed our model’s prediction that VANGL-mediated DVL phosphorylation plays a role in the PCP reaction network.

Mathematical modeling has historically proved a powerful approach for studying the biochemical network that underlies planar cell polarity. Perhaps the most well-known model is that by Amonlirdviman et al in 2005^69^ that used a simplified reaction framework to provide the first theoretical proof for how domineering non-autonomy phenotypes could arise purely from local interactions. While various models have been employed since, they generally fall into the category of either mass-action reaction-diffusion models^46,50^ or abstract physics-informed models^51,52^. Both have their drawbacks. On one hand, reaction-diffusion models can have large numbers of parameters and high computational requirements, making them challenging to scale beyond a few cells. On the other hand, abstract or phenomenological models, while useful for studying tissue scale polarity, often fail to tell us anything about the underlying mechanisms of PCP. We sought to balance these approaches by using a compartmental ODE model, which allows for a mechanistic reaction scheme while still maintaining the ability to model a large field of cells. Using this approach, we focused our model on key recent experimental findings, such as CK1-mediated VANGL and DVL phosphorylation^70^, allowing the model to offer a high level of mechanistic insight into PCP.

Whether or not cells can develop robust planar polarity alignment in the absence of an extrinsic instructional signal is an unresolved question. While WNT ligands sometimes provide such a signal in vivo^5^, in other cases polarity develops even with complete local knockout of WNT ligands^6,7^. Although it is impossible to truly prove the absence of a directional cue, our model supports a strong role for PCP-intrinsic alignment mechanisms in driving multicellular polarity alignment. Using only the biologically identified reaction schema, we show that positive feedback in the PCP pathway is sufficient to produce bistability at single membrane junctions, with the two stable states corresponding to the two potential orientations of the PCP complex. Furthermore, our multicellular modeling demonstrates that under the right conditions these bistable junctions are self-aligning, with neighboring junctions and cells aligning their direction of polarity due to simple diffusion. While this phenomenon is likely context dependent, our results demonstrate that intrinsic alignment mechanisms play a significant role in PCP biology, thus offering fresh insight into the spectrum of PCP phenotypes observed in different biological contexts.

Previous knockout experiments demonstrated that positive feedback through both VANGL and DVL are required to achieve robust polarity in vivo^31,71^ but our data also predict a unique role for each protein within the PCP feedback system. Specifically, our model predicts an important role for VANGL binding and clustering in the initial establishment of polarity, consistent with previous observations of dense puncta of PCP proteins in vivo^72^. This is likely due to the fast time scale of this feedback, as our analysis found that robust recruitment to the PCP complex and low dissociation rates from clustering puncta were key parameters for generating bistability (**Fig 2E)**. Conversely, we find that regulation of DVL phosphorylation was important for polarity alignment, as both robust phosphorylation at the membrane and rapid cytoplasmic de-phosphorylation were common in parameter sets that generated multicellular polarity alignment (**Fig 3I**). Interestingly, DVL is known to have many possible phosphorylation sites, only some of which are related to membrane retention and PCP^73^. Some sites, for example, promote the formation of cytoplasmic DVL puncta^73^, a process likely to sequester DVL and thus have the opposite effect on DVL membrane availability. Given these results, further study will be needed to understand the role of other DVL phosphorylation sites on PCP polarity.

An unexpected result from our model was the predicted role of VANGL phosphorylation in mediating the cellular response to WNT gradients. Previous studies have shown that a PCP-mediated phosphorylation cascade of VANGL at multiple sites near its N-terminus protects it both from recruitment into endocytic vesicles and from ubiquitin-mediated degradation^42,43^. Although the mechanisms underlying this behavior are not fully understood, our data predict that this differential stability is key to determining how cells polarize in response to a WNT gradient (**Fig 4**). Interestingly, WNT molecules are known to be alternatively chemoattractive^59^ or chemorepulsive^60^ to migrating cell collectives depending on context, a conundrum that has thus far had no mechanistic explanation. Our results predict that cellular differences in VANGL and pVANGL recycling are responsible for this difference, with robust endocytosis and degradation of VANGL predicted to be key to making WNT a repulsive ligand. Interestingly, this may have significant implications for cancer biology. Expression of non-canonical WNT ligands by tumors has been shown to be either oncogenic or tumor-suppressive, depending on the cancer type^74^. Furthermore, elevated expression of VANGL^18^, dysregulated endocytosis^75^, and dysregulated ubiquitination^76^ are all strongly associated with metastatic cancer, while knockout of VANGL2 drastically reduces tumor metastasis^18^. Our results provide a mechanistic link for these observations, suggesting that disruption of these systems has the potential to drastically alter planar cell polarity. Thus, targeting PCP may open the door to new lines of therapeutic research.

Recent studies support the existence of negative feedback loop in PCP whereby VANGL acts as a scaffold for DVL phosphorylation in the absence of FZD. Most notably, VANGL2, DVL, and CK1 co-IP as a single trimeric complex^38^, DVL is phosphorylated by CK1 in the absence of FZD binding^34^, and VANGL overexpression or knockdown directly correlate with pDVL levels^18^. Our modeling results demonstrate that addition of negative feedback enhances intracellular polarity alignment (**Fig. 3D**) and mediates the cellular response to WNT gradients (**Fig. 4E,F**), without significantly impairing the system’s inherent ability to polarize (**Fig. 2B,F**), while our experimental results support the existence of this mechanism through the observation of a clone-oriented migration phenotype (**Fig. 7**). Interestingly, this phenotype only occurred in a subset of simulations, indicating that it may be a context-dependent phenomenon. As only a handful of publications have investigated the VANGL overexpression domineering non-autonomy phenotype^31,77,78^, further research will be needed to fully understand the broad applicability and biological relevance of these findings.

Several studies have pointed toward important roles for planar cell polarity in the intestine in development, homeostasis, and disease. In development, PCP is critical for gut elongation^79^, the directed migration of sub-mucosal fibroblasts^80^, correct innervation of the intestinal tract by growing nerve fibers^81^, and Paneth cell differentiation^82^. In regenerative medicine, PCP-related non-canonical WNT ligands promote crypt fission following irradiation^83^, significantly reduce UC-related inflammation and relapse^15^, and impair tumor growth and metastasis in colorectal cancer^84^. Despite these important findings however, a direct mechanistic role for PCP in epithelial homeostatic migration or wound healing has not yet been established, although a previous report did demonstrate that intestinal epithelial cells undergo a remarkably uniform collective migration during homeostasis^23^. Our data using primary human intestinal epithelial cells show an important role for classical PCP signaling in the collective migration of both healthy, unperturbed hIEC monolayers (**Fig. 6**) and during the wound healing process (**Fig. 7**), shedding new light on the observed correlations between PCP dysregulation and intestinal disease, and establishing a new biological model system for studying PCP with strong clinical relevance.

Planar cell polarity is an essential biological process with an increasingly appreciated role in human disease. By combining computational modeling with a new biological platform to study PCP at a system level, our study provides new insights into the PCP feedback system and represents a critical step forward in our understanding of PCP biology.

## Methods

### General Approach

Our models utilize a compartmental ODE setup. That is, biological reactions within the model take place within well-mixed compartments, following mass action derived ordinary differential equations. Species concentrations at each time step are computed using the Euler method with dt=0.001. In the single junction model, each of the two cells are comprised of a single cytoplasmic compartment and a single membrane compartment. In the multicellular model, each cell is composed of a single shared cytoplasmic compartment and six membrane compartments, each of which forms an interaction junction with one of six neighboring cells. Cytoplasmic species (A, P, D, and Dp) shuttle between the cytoplasm and membrane compartments based on their on/off rates, and lateral diffusion between adjacent membrane compartments is calculated using the diffusion equation with dx=1 and Dc=1.75. CELSR dimers and any species bound to these dimers are assumed to not diffuse between adjacent compartments. Model simulations are generally initiated with all species having a starting concentration of 0.01 and iterated forward in time using the Euler method with dt=0.001. In multicellular simulations, the concentration of all cytoplasmic species in each cell are randomly initiated between 0 and 1. Gaussian white noise of strength D is added to the concentrations of each monomeric species at each time step to simulate intrinsic fluctuations in gene expression.

### Core Feedback Network

Our model is centered around the positive feedback loop we identify as a core mechanism of PCP, which we break down into six conceptual steps:

1. FZD proteins bind to CELSR dimers and induce low-level DVL phosphorylation
2. pDVL binding to the FZD-CELSR side of nascent PCP complexes stabilize them
3. Stabilized PCP complexes increase retention of VANGL-PRICKLE on the opposing membrane
4. Bound VANGL-PRICKLE initiates local clustering of VANGL, greatly stabilizing the complex
5. pDVL dissociates more readily than clustered VANGL, allowing the FZD side of the stabilized complex to become a robust phosphorylation site for additional pDVL
6. pDVL diffuses and stabilizes additional nascent PCP complexes

The core inhibitory mechanism is generated by the phosphorylation of VANGL at the FZD-CELSR side of PCP complexes. This phosphorylation inhibits both VANGL binding and clustering to opposite orientation nascent clusters, breaking the opposite-orientation positive feedback loop.

### Additional Reactions

In addition to this core feedback network, several other mechanisms have been identified to regulate PCP and were included in our modeling. These include: **1)** diffusive membrane species bound to PRICKLE demonstrate enhanced endocytosis and degradation, **2)** VANGL enhances PRICKLE and DVL cytoplasm-to-membrane recruitment in a concentration-dependent manner, **3)** DVL phosphorylation allows it to dimerize, further enhancing its membrane stability and complex-stabilizing effect, and **4)** PRICKLE competes with ANKRD6 in binding to non-phosphorylated DVL to inhibit its phosphorylation.

### Key Model Assumptions

To ensure our models of the PCP reaction network were computationally feasible, several key assumptions were made regarding the biochemical interactions involved. **1)** Given a lack of evidence otherwise, we assumed that the binding and dissociation of secondary cytoplasmic proteins (ANKRD6/PRICKLE) to their key binding partners (DVL/VANGL) are independent of whether these proteins are in complex with CELSR or FZD. **2)** As cellular regulation of CK1 concentration has not been shown to be a key regulatory mechanism of PCP, the concentration of the kinase CK1 was assumed to be non-limiting. **3)** While the exact role of ANKRD6 is not well studied in the context of PCP, early studies indicated it has a key role in recruiting the kinase CK1^85^, thus phosphorylation of DVL was modeled to be dependent on the presence of bound ANKRD6. **4)** Similarly, while the exact role of PRICKLE in the PCP reaction network is not well known, its essential requirement in polarity establishment and the presence of four protein interaction domains (1 PET and 3 LIM domains in PRICKLE1/2) has led to the supposition that is a key regulator of VANGL clustering^86^. Thus, we assume VANGL clustering to be dependent on the concentration of PRICKLE bound to fully formed PCP complexes. **5)** We assume that PCP membrane phosphorylation dynamics occur on a much slower time scale than cytoplasmic diffusion and thus do not model cytoplasmic diffusion (i.e. we assume a well-mixed cytoplasm at all time steps). **6)** We do not include in our model any stable cytoplasmic complexes. While complexes such as cytoplasmic DVL aggregates do exist^87^, we assume these form an additional level of regulation outside of the scope of our current models. **7)** We do not model any directed transport of PCP proteins. While directed vesicle trafficking of PCP proteins along microtubules has been previously shown^2^, its contribution to PCP polarity is unknown, and we find it unnecessary to generate experimental phenotypes. **8)** Given a lack of evidence otherwise, membrane proteins were modeled to diffuse between membrane compartments at the same rate. **9)** We do not model differences between protein isoforms.

### Parameter Space Boundaries and Model Constraints

In general, most parameter rates were bounded to be within two orders of magnitude, between 0.1 and 10. Notable exceptions were k1 and k2, the binding and dissociation rates for CELSR dimers, as variation in these rates was found drive wildly different complex concentrations. To ensure similar PCP complex concentrations across parameter sets, these parameters were bounded between 0.05 and 0.2. Synthesis and degradation rates were assumed to occur on a slower timescale than membrane interactions and were also bounded between 0.05 and 0.2.

Beyond these bounds, additional biologically informed constraints were placed upon the parameter sets used to reduce the overall size of parameter space. Specifically, **1)** DVL phosphorylation and dimerization increases the membrane on rate and decreases the membrane off rate (D_on_ < Dp_on_) and (D_off_ > Dp_off_ > DpDp_off_). **2)** DVL phosphorylation and dimerization increases its binding rate to FZD and reduces its dissociation rate (k25a < k25b < k25c) and (k26a > k26b > k26c). **3)** The binding of DVL, pDVL, and dimerized pDVL to FZD successively decreases the rate at which FZD dissociates from CELSR dimers (k4a < k4b < k4c < k4d). **4)** VANGL phosphorylation increases its basal dissociation rates from PCP complexes and reduces the protection offered by VANGL clustering (k7min < k8min), (k7m < k8m), and (k7max < k8max). **5)** VANGL phosphorylation reduces its incorporation into VANGL clusters (k27a > k27b) and (k24a < k24b). **6)** Membrane proteins in complex with PRICKLE, but not bound to CELSR dimers, have higher degradation rates than the unbound form of the species (dVP > dV), (dVpP > dVp), and (dFDP > dF).

### Primary hISC culture and transfection

Primary human intestinal stem cell (hISC) culture was carried out using published methods^62^, in which stem cells are maintained in two-dimensional culture on a thick layer of collagen. Briefly, aqueous acidic Rat Tail Collagen I (R&D Systems #3443-100-01) is neutralized with a sodium hydroxide solution in standard tissue culture plates to induce the formation of a thick 1 mg/mL collagen gel. This gel is rinsed three times with PBS, then primary intestinal crypts or dissociated cells are seeded directly on top. Culture media also follows previously published methods^63^ consisting of 50% WNR (Wnt, Noggin, Rspondin) conditioned media, diluted with 2x Basal medium comprised of Advanced DMEM/F12 (Gibco #12634010) containing: 4% B27 without Vitamin A (Gibco #12587001), 10 mM Nicotinamide (Sigma-Aldrich N0636), 20 mM HEPES (Corning #25-060-Cl), 1X Glutamax (Gibco #35050061), 1X Pen/Strep (Gibco #15070063), 1.25 mM N-Acetylcysteine (Sigma-Aldrich #A9165), 50 ug/mL Primocin (Invivogen #ant-pm-05), 3 uM SB202190 (Peprotech #1523072), 50 ng/mL mEGF (Peprotech #315-09), 2 nM Gastrin (Sigma-Aldrich #G9145), and 10 nM Prostaglandin E2, (Peprotech #3632464).

Primary hISCs were isolated as previously described^63^. Briefly, anonymous intestinal tissue samples were obtained from cadaveric organ donors undergoing surgical organ donation. Stem cell rich epithelial crypts from the mid-jejunum were isolated by repeated agitation in an EDTA-DTT solution, then plated onto collagen-coated plates in stem cell maintenance media plus Y27 and grown in a standard tissue culture incubator at 37 C, 5% CO2. Media was replaced the following day with Y27-free medium and fed every other day thereafter. After 5-7 days, and every 5-7 days thereafter when the cells reached ∼80% confluency, the cells were passaged at a 1:3 split ratio. To passage, the collagen patty was mechanically released from the plate and digested with collagenase IV (750 U/mL final concentration) at 37 C for 20 minutes (Worthington Biochemical #LS004188). The resulting cells were rinsed with PBS, incubated in warm TrypLE Express (ThermoFisher Scientific #12605010) for 5 minutes at 37 C, then mechanically dissociated by pipet trituration. TrypLE was quenched with media, and the dissociated cells were pelleted and resuspended in fresh stem cell maintenance media plus Y27, plating onto new collagen-coated plates. Media was replaced the day after passage and every other and every other day thereafter.

To generate the WT^H2B-GFP^ and VANGL2-iOE^H2B-RFP^ polyclonal cell lines, primary hISCs below passage 5 were transfected following previously optimized methods^63^, using the Neon electroporation system to co-transfect Supper PiggyBAC Transposase™ expression plasmid alongside ITR-containing payload plasmids. Cells were recovered in culture for 3 days, then antibiotic selection (2 ug/mL Puromycin for WT^H2B-GFP^ or 6.67 ug/mL Blasticidin for VANGL2-iOE^H2B-GFP^) was applied for five days. The resulting polyclonal cell populations were expanded, and transgene integration was confirmed by long-term expression of the H2B fluorescent tags in >95% of the cells.

### Preparation of Planar Crypt Microarray Transwell Inserts

PET film with laser-drilled microhole patterns were obtained from ALTIS Biosystems, washed with a 1% Tergezyme (MilliporeSigma #Z742918) solution for 5 minutes, then rinsed thoroughly with DI water. The permeable membranes of commercial 12-well transwell inserts (Corning #3401) were removed with forceps, and the PCM membranes were adhered using Loctite 3979 UV-cured acrylic adhesive gel, cured for 2 minutes under a 50W 365 nm UV light source. A positive charge was applied to the membrane with a vacuum oxygen plasma treatment, 20W for 1 minute. Membranes were pre-warmed to 37 C. Rat tail collagen was neutralized on ice, and 200 uL of neutralized collagen was added to each well and polymerized for 1 hour at 37 C. For wound assay experiments, a thin strip (0.8 mm x 20 mm) of plasma-treated PET film, dubbed a “wound strip”, was bent into an L shape, and the lower half of the L shape was dipped into freshly neutralized ice-cold 1 mg/mL collagen, and placed on top of the first collagen layer just outside of the microhole region, followed by a second 1 hour 37 C incubation. Following collagen polymerization, PCMs were transferred to a 40 C dry oven and incubated overnight (∼16 hours), during which the collagen dried to form a compact layer. Dried PCMs were stored at room temperature until use.

### Cell Culture and Live Imaging Assays on PCMs

Before plating, dried PCMs were rinsed for 1 minute in DI water to remove excess salt, then sterilized in a 70% ethanol bath for 5 minutes. Sterile PCMs were rinsed three times with excess PBS, then 500 uL of 1% Matrigel in PBS was added to the apical compartment and incubated for at least 1 hour at 37 C. After 1 hour the Matrigel solution was aspirated, and PCMs were rinsed twice with PBS.

Primary hISCs were dissociated and resuspended in fresh media plus Y27 as above and 500 uL of cell suspension was plated on the apical side of PCMs at a split ratio of 1:4 (ie. 1x well of a 6- well plate to 4x PCMs on 12-well Transwell inserts. 1 mL of warmed MM+Y was added to the basal compartment of each PCM, and cells recovered overnight in the incubator. Media was replaced the following day with fresh pre-warmed MM, and cells were visually inspected to ensure they had formed a confluent monolayer over the entire PCM growth area. PCMs were fed daily with fresh MM (500 uL MM apical, 1 mL MM basal) until initial compartmentalization was observed over the micro- holes using a standard inverted phase-contrast microscope, typically 5 days post-plating, at which point apical media was replaced with 500 uL Differentiation Media, comprised of Advanced DMEM/F12 (Gibco #12634010) containing: 10 mM HEPES (Corning #25-060-Cl), 1X Glutamax (Gibco #35050061), 1X Pen/Strep (Gibco #15070063), 1.25 mM N-Acetylcysteine (Sigma-Aldrich #A9165), 50 ug/mL Primocin (Invivogen #ant-pm-05), 50 ng/mL mEGF (Peprotech #315-09), and 500 uM A83- 01, (Sigma-Aldrich #SML0788). The basal compartment was fed with 1 mL MM as before (DM/MM). At 24 hours of differentiation, media was replaced with DM/MM +/- 500 ng/mL doxycycline hyclate (Sigma-Aldrich #D9891-25G). At 48 and 64 hours, apical media was replaced with 500 uL DM mixed 1:1 with Wnt5a-conditioned media, feeding with standard MM basally.

Live imaging was initiated after 72 hours of differentiation, and most imaging was performed on Keyence BZ-X810 inverted fluorescence microscope combined with a Tokai Hit STRF-KIW live imaging chamber. Confocal imaging was carried out using an Andor Dragonfly spinning disk confocal microscope with an iXon Life 888 EMCCD camera. In both cases, images were taken every 15 - 30 minutes for at least 12 hours, and all images being directly compared were captured using identical settings. For wound assay experiments, wounds were induced immediately prior to imaging by using a pair of forceps to grab the protruding segment of the wound strip and pulling gently, tearing up a thin strip of cells.

### Immunofluorescent Staining

For experiments with planned immunofluorecent staining, media was aspirated and PCMs were fixed in 4% PFA (500 uL apical, 500 uL basal, ThermoFisher Scientific #AC416780250) for 20 minutes at room temperature, followed by 3x rinses with excess PBS. Cells were permeabilized with 0.5% Triton X-100 (ThermoFisher Scientific AC215682500) for 20 minutes, then rinsed three times with PBS+3% BSA (FisherScientific BP1600-1). Blocking was performed for two hours at room temperature in 3% BSA. After blocking, cells were incubated with either sheep anti-VANGL2 (1:100 dilution, R&D Systems #AF4815) or rabbit anti-FZD6 (1:100 dilution, Antibodies.com #A99895) primary antibody in 3% BSA overnight (16 hours) at 4°C, rinsed three times with IF wash solution (0.1% BSA, 0.2% Triton-X 100, and 0.05% Tween-20 in PBS, 5 minute incubation per rinse), then incubated with donkey anti-sheep Alexa Fluor™ 488 (1:500, Abcam #ab150177) or donkey anti-rabbit Alexa Fluor™ 488 (1:500, Abcam #ab150073) secondary antibody and DAPI (200ng/mL final concentration, ThermoFisher #D3571) in 3% BSA for 1 h at room temperature. Stained cells were rinsed three times with 3% BSA, then stored in 3% BSA for microscopic analysis.

### Cell Tracking of Live Imaging Experiments

To track homeostatic cell migration on PCMs, the Manual Tracking plugin in Image J was used to record cell positions at each time point. Forty-eight tracks were recorded per replicate for a total of ∼144 tracks per condition, following cells that begin the time-course at least 50 uM from any microhole. The net displacement vector (𝑥𝑦_𝑓𝑖𝑛𝑎𝑙_ − 𝑥𝑦_𝑖𝑛𝑖𝑡𝑖𝑎𝑙_) was calculated for each track, and the net displacement angle was calculated as the difference between the angle of the net displacement vector and a radial vector extending from the PCM center and intersecting 𝑥𝑦_𝑖𝑛𝑖𝑡𝑖𝑎𝑙_. For the full python code used in this analysis, see Data and Code Availability. For high throughput tracking during wound healing, the Imaris Imaging Suite spot tracking feature was used with the built-in autoregressive motion tracking algorithm, filtering out tracks lasting less than 10 hours.

## Acknowledgements

We would especially like to thank the anonymous organ donors and their families for their contributions, as well as HonorBridge organ procurement organization for providing tissue samples used in this study. Figure schematics were created with BioRender.com. The Keyence microscope was funded with support from CGIBD core grant P30DK034987. The Andor Dragonfly microscope was funded with support from National Institutes of Health grant S10OD030223. This work was funded by NIH grants T32GM133364 (K.A.B.), 5F31DK136305 (K.A.B.), R35GM127145 (T.C.E.), R01DK115806 (S.T.M.), and R01DK109559 (S.T.M.). PCM membranes were obtained from Altis Biosciences, in which Scott Magness has a financial interest.

