## Supplemental Data for "Coupling mathematical modeling with a novel human intestinal stem cell system to understand feedback regulation during planar cell polarity"

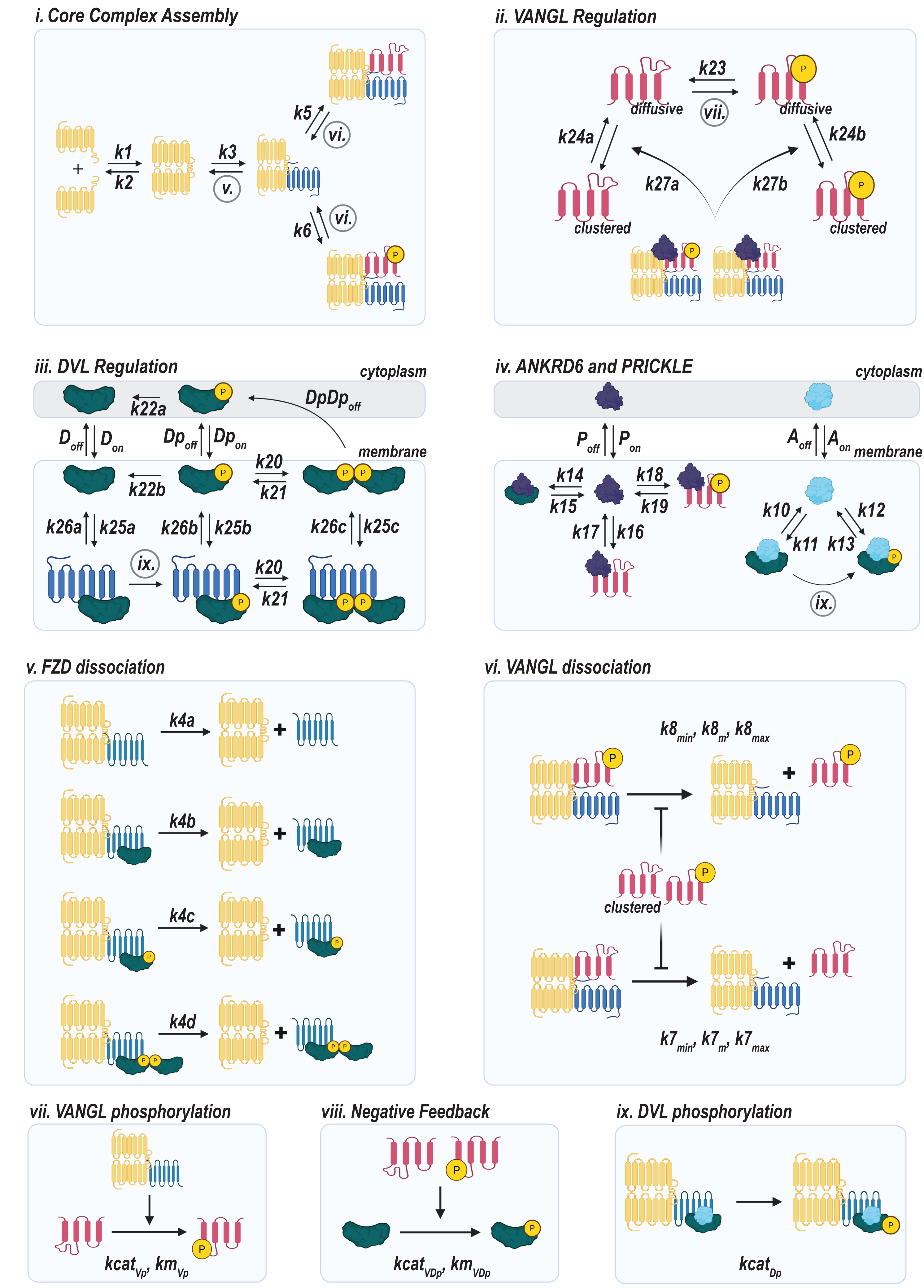


### Figure S1 - Graphical Overview of Reaction Network

A

##### Sythesis and Degredation

| Parameter | Min Value | Max Value | Description |
|---|---|---|---|
| sD | 0.05 | 0.2 | DVL synthesis |
| sA | 0.05 | 0.2 | ANKRD6 synthesis |
| sP | 0.05 | 0.2 | PRICKLE synthesis |
| sV | 0.05 | 0.2 | VANGL synthesis |
| sF | 0.05 | 0.2 | FZD synthesis |
| sC | 0.05 | 0.2 | CELSR synthesis |
| dD | 0.05 | 0.2 | DVL degredation |
| dA | 0.05 | 0.2 | ANKRD6 degredation |
| dP | 0.05 | 0.2 | PRICKLE degredation |
| dC | 0.05 | 0.2 | CELSR degredation |
| dV | 0.05 | 0.2 | VANGL degredation |
| dVP | 0.1 | 0.5 | VANGL degredation when bound to PRICKLE |
| dVp | 0.05 | 0.2 | Phospho-VANGL degredation |
| dVpP | 0.1 | 0.5 | Phospho-VANGL degredation when bound to PRICKLE |
| dF | 0.05 | 0.2 | FZD degredation |
| dFDP | 0.1 | 0.5 | FZD degrdation when bound to DVL-PRICKLE |
| k9a | 0.1 | 10 | V-dependent degredation of P |
| k9b | 0.1 | 10 | Vp-dependent degredation of P |
| k9c | 0.1 | 10 | Dp-dependent degredation of P |

##### Core Complex Assembly

| Parameter | Min Value | Max Value | Description |
|---|---|---|---|
| k1 | 0.05 | 0.2 | CELSR Dimerization |
| k2 | 0.05 | 0.2 | CELSR De-dimerization |
| k3 | 0.2 | 10 | Binding of FZD to CELSR dimer |
| k4a | 0.1 | 10 | Unbinding of FZD from CELSR dimer |
| k4b | 0.1 | 10 | Unbinding of FZD-DVL from CELSR dimer |
| k4c | 0.1 | 10 | Unbinding of FZD-pDVL (monomer) from CELSR dimer |
| k4d | 0.1 | 10 | Unbinding of FZD-pDVL (dimerized) from CELSR dimer |

##### ANKRD6 and PRICKLE

| Parameter | Min Value | Max Value | Description |
|---|---|---|---|
| k10 | 0.1 | 10 | ANKRD6 binding to DVL |
| k11 | 0.1 | 10 | ANKRD6 dissociation from DVL |
| k12 | 0.1 | 10 | ANKRD6 binding to pDVL |
| k13 | 0.1 | 10 | ANKRD6 dissociation from pDVL |
| k14 | 0.1 | 10 | PRICKLE binding to DVL |
| k15 | 0.1 | 10 | PRICKLE dissociation from DVL |
| k16 | 0.1 | 10 | PRICKLE binding to VANGL |
| k17 | 0.1 | 10 | PRICKLE dissociation from VANGL |
| k18 | 0.1 | 10 | PRICKLE binding to phospho-VANGL |
| k19 | 0.1 | 10 | PRICKLE dissociation from phospho-VANGL |

##### Membrane On/Off Rates

| Parameter | Min Value | Max Value | Description |
|---|---|---|---|
| D_on | 0.1 | 10 | DVL membrane association |
| D_off | 0.1 | 10 | DVL membrane dissociation |
| DP_off | 0.1 | 10 | DVL membrane dissociation when bound to PRICKLE |
| Dp_on | 0.1 | 10 | Phospho-DVL membrane association |
| Dp_off | 0.1 | 10 | Phospho-DVL membrane dissociation |
| DpDp_off | 0.1 | 10 | Dimerized phospho-DVL membrane dissociation |
| A_on | 0.1 | 10 | ANKRD6 membrane association |
| A_off | 0.1 | 10 | ANKRD6 membrane dissociation |
| P_on | 0.1 | 10 | PRICKLE membrane association |
| P_off | 0.1 | 10 | PRICKLE membrane dissociation |
| PV_on | 0.1 | 10 | VANGL-dependent recruitment of P to the membrane |
| DV_on | 0.1 | 10 | VANGL-dependent recruitment of D to the membrane |

##### VANGL Regulation

| Parameter | Min Value | Max Value | Description |
|---|---|---|---|
| k5 | 0.1 | 10 | Binding of VANGL to CELSR-FZD complexes |
| k6 | 0.1 | 10 | Binding of pVANGL to CELSR-FZD complexes |
| k7min | 0.1 | 10 | Min dissociation of VANGL from CELSR-FZD complexes |
| k7m | 0.1 | 10 | Half-max dissociation of VANGL from CELSR-FZD complexes |
| k7max | 0.1 | 10 | Max dissocaition of VANGL from CELSR-FZD complexes |
| k8min | 0.1 | 10 | Min dissociation of pVANGL from CELSR-FZD complexes |
| k8m | 0.1 | 10 | Half-max dissociation of pVANGL from CELSR-FZD complexes |
| k8max | 0.1 | 10 | Max dissocaition of pVANGL from CELSR-FZD complexes |
| k23 | 0.1 | 10 | VANGL de-phosphorylation |
| k24a | 0.1 | 10 | VANGL de-clustering |
| k24b | 0.1 | 10 | Phospho-VANGL de-clustering |
| k27a | 0.1 | 10 | VANGL clustering |
| k27b | 0.1 | 10 | Phospho-VANGL clustering |
| kcat_Vp | 0.1 | 10 | Catalytic rate of VANGL phosphorylation by FZD |
| km_Vp | 0.1 | 10 | Michaelis-Menton phosphorylation of Vp half-max |

##### DVL Regulation

| Parameter | Min Value | Max Value | Description |
|---|---|---|---|
| k20 | 0.1 | 10 | pDVL Dimerization |
| k21 | 0.1 | 10 | pDVL Un-dimerization |
| k22a | 0.1 | 10 | Cyotoplasmic de-phosphorylation |
| k22b | 0.1 | 10 | Membrane de-phosphorylation |
| kcat_Dp | 0.1 | 10 | DVL phosphorylation by FZD |
| kcat_VDp | 0.1 | 10 | DVL phosphorylation by VANGL |
| km_VDp | 0.1 | 10 | DVL phosphorylation by VANGL half-max |
| k25a | 0.1 | 10 | DVL association with FZD |
| k25b | 0.1 | 10 | Phospho-DVL (monomer) association with FZD |
| k25c | 0.1 | 10 | Phospho-DVL (dimerized) association with FZD |
| k26a | 0.1 | 10 | DVL dissociation from FZD |
| k26b | 0.1 | 10 | Phospho-DVL (monomer) dissociation from FZD |
| k26c | 0.1 | 10 | Phospho-DVL (dimerized) dissociation from FZD |

B

| Constraint | Description |
|---|---|
| D_on < Dp_on | Phosphorylation increases DVL membrane recruitment |
| D_off > Dp_off > DpDp_off | Phosphorylation decreases DVL membrane dissociation |
| k25a < k25b < k25c | DVL association with FZD increases with phosphorylation |
| k26a > k26b > k26c | DVL dissociation from FZD decreases with phosphorylation |
| k4a > k4b > k4c > k4d | Dissociation of FZD from CELSR decreases with D, Dp, and DpDp bound |
| k7min < k8min | Basal dissociation of VANGL from PCP complexes increases with phorphorylation |
| k7m < k8m | Amount of clustered VANGL required to inhibit dissocaition increases with phosphorylation |
| k7max < k8max | Max dissociation rate of VANGL from PCP complexes increases with phorphorylation |
| k27a > k27b | VANGL clustering decerases with phosphorylation |
| k24a < k24b | De-clustering of VANGL increases with phosphorylation |
| dV < dVP | PK binding increases VANGL degredation rate |
| dVp < dVpP | PK binding increases phospho-VANGL degredation rate |
| dF < dFDP | PK binding increases FZD degredation rate |

### Figure S2 - Parameter and Constraint Tables

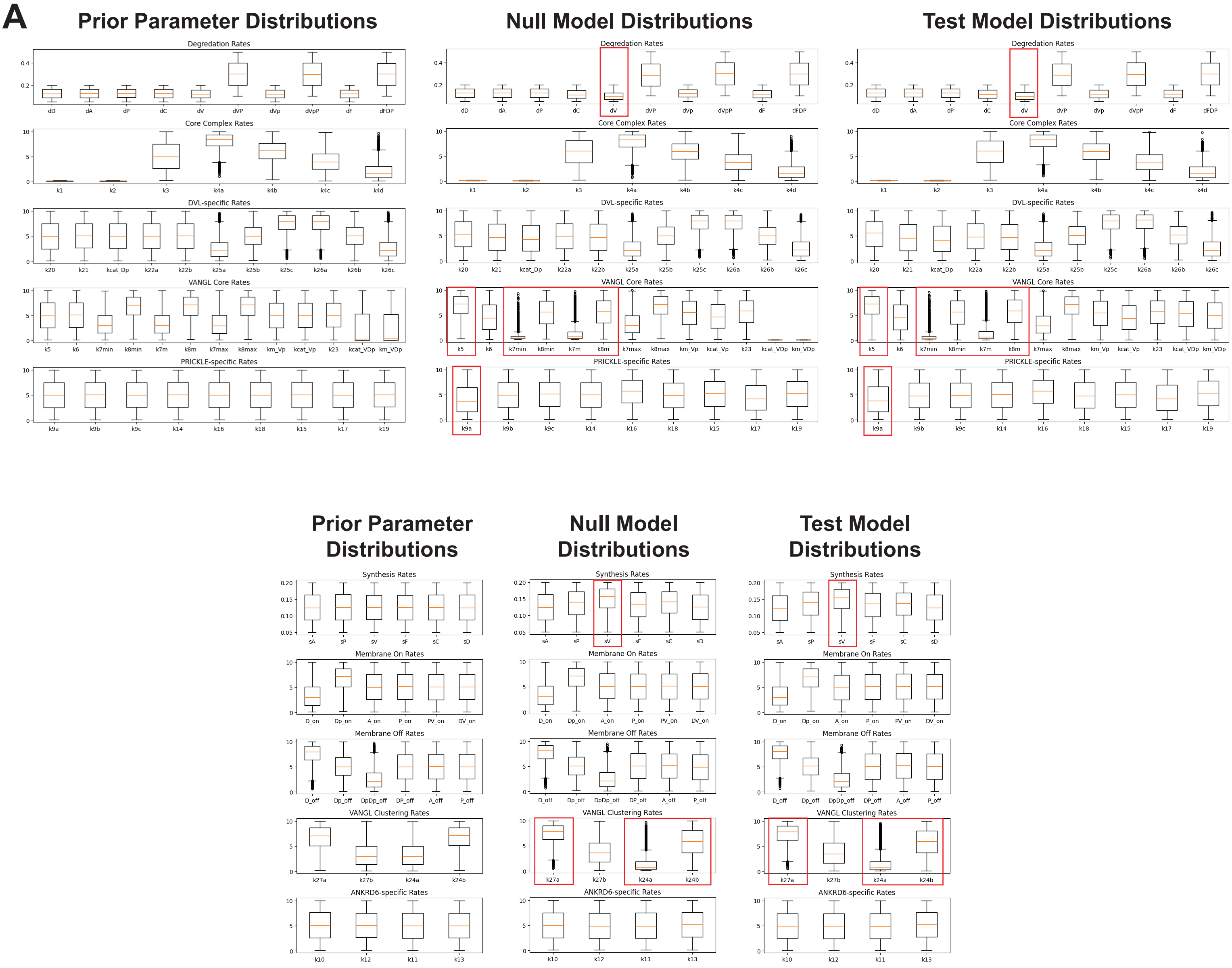


### Figure S3 - Bistable Parameter Distributions

**Figure S1. Graphical Overview of Reaction Network**

Overview of the model reaction scheme employed showing all component interactions and labeled with the rate constants involved. Circled roman numerals indicate a more complicated reaction than what is shown in that panel and reference the panel in which the full reaction can be found.

**Figure S2. Parameter and Constraint Tables**

**(A)** Parameter names, bounds, and descriptions for all parameters used in the model, separated into categories based on function. **(B)** Constraints placed upon initial parameter values to reduce the overall size of parameter space.

**Figure S3. Bistable Parameter Distributions**

Parameter distributions of the initial 5 million parameter sets (Prior) compared to the parameter distributions observed in the subsets of bistable parameter sets for the Null and Test models. Red boxes indicate the top 11 most significant parameters contributing to bistability, all of which relate to VANGL.
